# Mapping human microglial morphological diversity via handcrafted and deep learning-derived image features

**DOI:** 10.1101/2025.09.12.675829

**Authors:** Kayhan Alvandipour, Amélie Weiss, Mona Mathews, Brenda Besemer, Michaela Segschneider, Zahra Hanifehlou, Michael Peitz, Arnaud Ogier, Peter Sommer, Oliver Brüstle, Johannes H. Wilbertz

**Affiliations:** Ksilink, Strasbourg, France; LIFE & BRAIN GmbH, Cellomics Unit, Bonn, Germany; Institute of Reconstructive Neurobiology, University of Bonn Medical Faculty & University Hospital Bonn, Bonn, Germany

## Abstract

Microglia play critical roles in brain health and disease by adopting a spectrum of dynamic activation states. However, capturing this continuous heterogeneity in a scalable way remains a major challenge. To address this, we developed an imaging and data analysis framework to map the activation landscape of human iPSC-derived microglia (iMG) at single-cell resolution. We combined high-content imaging using two complementary strategies: a hypothesis-driven immunofluorescence (IF) panel targeting key activation markers (NF-κB, ASC, CD45) and a discovery-oriented, pan-morphological Cell Painting (CP) assay. Diverse phenotypes were captured through handcrafted and representation learning-based features. To classify cells, we applied Gaussian Mixture Models (GMMs) to image-derived features, enabling soft probabilistic assignments that capture transitional states between phenotypes. Compared to graph-based methods like the Leiden algorithm, GMMs provided comparable classification performance while offering a more nuanced and biologically interpretable view of microglial heterogeneity. We demonstrate that deep learning features from the targeted IF panel are most powerful, achieving high classification accuracy and strong correlation with biological states such as functional NLRP3 inflammasome activation. Our model system provides a robust and scalable platform for quantifying microglial heterogeneity, offering a new tool to identify novel disease-associated states and compounds that precisely modulate microglial phenotypes for therapeutic discovery.

## Introduction

Microglia, the resident immune cells of the central nervous system, exhibit highly dynamic and context-dependent phenotypes in response to inflammatory stimuli ^1^. Accurate phenotyping of their activation states is essential for understanding their roles in neuroinflammation and neurodegeneration. However, no standardized, high-throughput-compatible methods currently exist for comprehensive microglial profiling. Resolving distinct subpopulations is also critical for therapeutic development, as targeting specific subsets could enable more precise and effective interventions. In contrast, commonly used CSF1R inhibitors act broadly on virtually all microglia and most tissue-resident macrophages by blocking a fundamental survival signals shared across these cell types. While this makes CSF1R inhibitors powerful tools for depleting microglia, their widespread effects complicate interpretation and limit their utility for selectively modulating disease-relevant microglial states ^2–4^.

Single-cell RNA sequencing (scRNA-seq) has been instrumental in uncovering microglial heterogeneity, identifying distinct subpopulations with specialized transcriptional signatures, including homeostatic microglia, disease-associated microglia (DAM), and age-specific phenotypes ^3–9^. These insights have deepened our understanding of microglial roles across development, brain regions, and disease contexts. However, transcriptomic data alone may not fully capture functional states, particularly those shaped by post-transcriptional regulation, protein localization, or morphological dynamics ^7,8,10,11^.

Imaging-based cellular phenotyping offers a powerful and complementary approach by providing direct, functional readouts at single-cell resolution. It leverages advanced microscopy and image analysis to quantify morphological, spatial, and molecular features of individual cells, enabling classification of cellular states and functions *in vitro* ^12,13^. Imaging is inherently scalable and compatible with high-throughput workflows, making it well-suited for large-scale phenotypic profiling. Yet, phenotypic characterization remains heavily dependent on the choice of staining markers and image analysis strategies, which can introduce bias and limit reproducibility.

To address this, we systematically characterized human induced pluripotent stem cell (iPSC)-derived microglia (iMG) ^14^ that were primed using lipopolysaccharide (LPS) and activated using Nigericin to induce the NLRP3 inflammasome, a multi-protein complex that detects cellular stress signals and activates inflammatory responses by promoting the maturation of cytokines such as IL-1β and IL-18. LPS binds to Toll-like receptor 4 (TLR4) on the microglial surface, initiating a signaling cascade that activates NF-κB. This priming involves the nuclear translocation of NF-κB, where it promotes the transcription of pro-inflammatory cytokines and key inflammasome components such as NLRP3 ^15,16^. Nigericin, a potassium ionophore, subsequently triggers the assembly of the NLRP3 inflammasome by inducing potassium efflux, a critical signal for inflammasome activation and caspase-1–mediated maturation of IL-1β ^17,18^.

To phenotypically characterize iMG we used two biologically relevant staining approaches: a targeted immunofluorescence (IF) panel focused on key activation markers, and a holistic Cell Painting (CP) assay designed to capture broad morphological diversity ^19^. These were paired with two distinct image analysis pipelines: classical handcrafted (HC) feature extraction and deep learning–based (DL) representation learning. Together, these approaches generated rich, multi-dimensional feature sets for single-cell clustering. However, a major bottleneck in imaging-based phenotyping is the lack of computational methods that can robustly cluster the continuous, dense data from cellular images to reveal transitional states without imposing artificial hard boundaries. To overcome the limitations of clustering algorithms commonly used in scRNA-seq analysis, such as Louvain, Leiden, and Phenograph, which are optimized for high-dimensional transcriptomic data and rely on graph-based representations, we applied a Gaussian Mixture Model (GMM) trained on the diverse imaging-derived feature sets generated in this study ^20,21^. By comparing GMM clustering to Leiden algorithm-based clustering, we show that GMMs are well-suited for continuous, dense, and morphologically structured data, enabling probabilistic modeling of phenotypic states and capturing subtle variations in cell morphology and marker expression. GMM clustering enables soft clustering, assigning cells to multiple clusters with varying degrees of membership, thereby offering a more nuanced and biologically interpretable view of microglial heterogeneity.

All experimental data used to train and validate the algorithms were generated from three fully independent iMG differentiations using a developmentally informed protocol ^14^ and separate imaging runs, conducted in 384-well plates. Our study design and methodological framework enable robust, scalable, and biologically diverse profiling of microglial activation dynamics at single-cell resolution, demonstrating strong generalizability across biological and technical variability inherent to iPSC-derived microglia models.

## Results

### Immunofluorescence and Cell Painting staining across iMG activation states

To investigate human microglial activation dynamics, we employed iMG generated from three independent differentiation batches (LB:MiGs; batches P1-P3). Cells were seeded in 384-well plates at 8,000 cells/well and stimulated using a two-step protocol: priming with LPS (4 hours) followed by activation with Nigericin (3 hours) (**Figure 1A**). Post-stimulation, cells were fixed and analyzed using two complementary imaging modalities. First, immunofluorescence (IF) staining targeted DNA, NF-κB, CD45, and ASC to assess nuclear translocation (NF-κB), cell identity/size (CD45), and inflammasome activation (ASC specks) (**Figure 1B**). Second, Cell Painting (CP), a high-content morphological profiling method, labeled DNA, mitochondria, RNA, endoplasmic reticulum (ER), and a composite channel for actin, Golgi, and plasma membrane (AGP) (**Figure 1C**). While CP is not microglia-specific, it enables broad phenotypic characterization.

**Figure 1.**
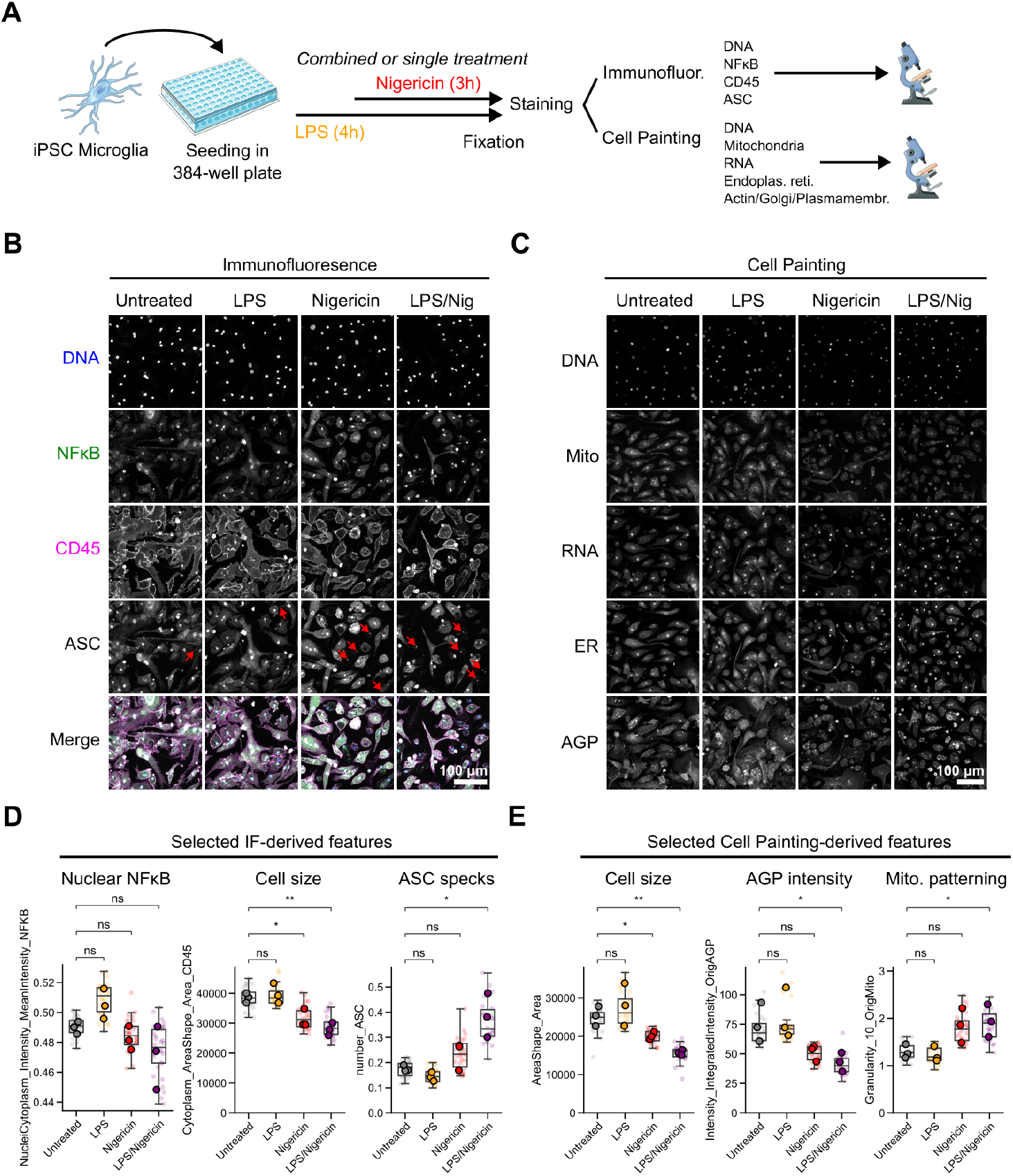
Experimental design and imaging-based profiling of iPSC-derived microglia. **(A)** Schematic overview of the stimulation protocol. iPSC-derived microglia were seeded in 384-well plates and treated with LPS (4 h), Nigericin (3 h), or their combination, followed by fixation and staining for either immunofluorescence (IF) or Cell Painting (CP). **(B)** Representative IF images showing staining for DNA (blue), NF-κB (green), CD45 (magenta), and ASC specks (red arrows) across treatment conditions. **(C)** Representative CP images highlighting cellular compartments stained for DNA, mitochondria (Mito), RNA, endoplasmic reticulum (ER), and actin/Golgi/plasma membrane (AGP). **(D)** Quantitative features extracted from IF images, including nuclear NF-κB intensity, cell size, and ASC speck count. **(E)** Quantitative features extracted from CP images, including cell size, AGP intensity, and mitochondrial texture. Boxplots show the distribution of technical replicates (mean of 9 images per 384-well plate well), with small markers for individual wells and larger markers for the means of three independent iMG differentiations. Statistical comparisons were performed using two-sided independent t-tests on biological replicate means. Boxplots display the median, interquartile range (IQR), and whiskers extending to 1.5×IQR.

Initial quantitative analysis focused on three IF-derived features: nuclear NF-κB intensity, CD45-based cell area, and ASC specks per cell (**Figure 1D**). LPS priming increased nuclear NF-κB, commonly observed in primed microglia ^15,16^. Nigericin treatment, alone or with LPS, resulted in a significantly reduced cell area, indicated by a compact, morphology reminiscent of activated cells ^1,22^. This reduction was corroborated by CP-derived measurements using cytoplasmic and membrane-associated stains (**Figure 1E**).

ASC speck formation, a confirmatory hallmark of canonical inflammasome assembly, was elevated after LPS/Nigericin activation (**Figure 1D**). CP features, including actin/Golgi/PM intensity and mitochondrial texture, also changed upon stimulation, reflecting cytoskeletal remodeling and metabolic reprogramming which are all hallmarks of microglial activation (**Figure 1E**). These findings validate the LPS/Nigericin priming/activation paradigm in iMG and highlight the complementary strengths of IF and CP for comprehensive microglial phenotyping.

### Single-cell phenotypic features correlate with iMG activation states

To systematically investigate iMG responses under varying inflammatory conditions, we employed a high-throughput imaging approach using a 384-well plate format and three independent differentiation batches, totaling 240 wells. Cells from all treatment conditions were imaged across 16 fields per well using IF and CP staining. This yielded 4,320 IF and 5,400 CP images, corresponding to approximately 62,921 and 59,260 single cells, respectively. These images formed the basis for downstream single-cell analysis, enabling robust phenotypic profiling across diverse activation states (**Figure 2A**).

**Figure 2.**
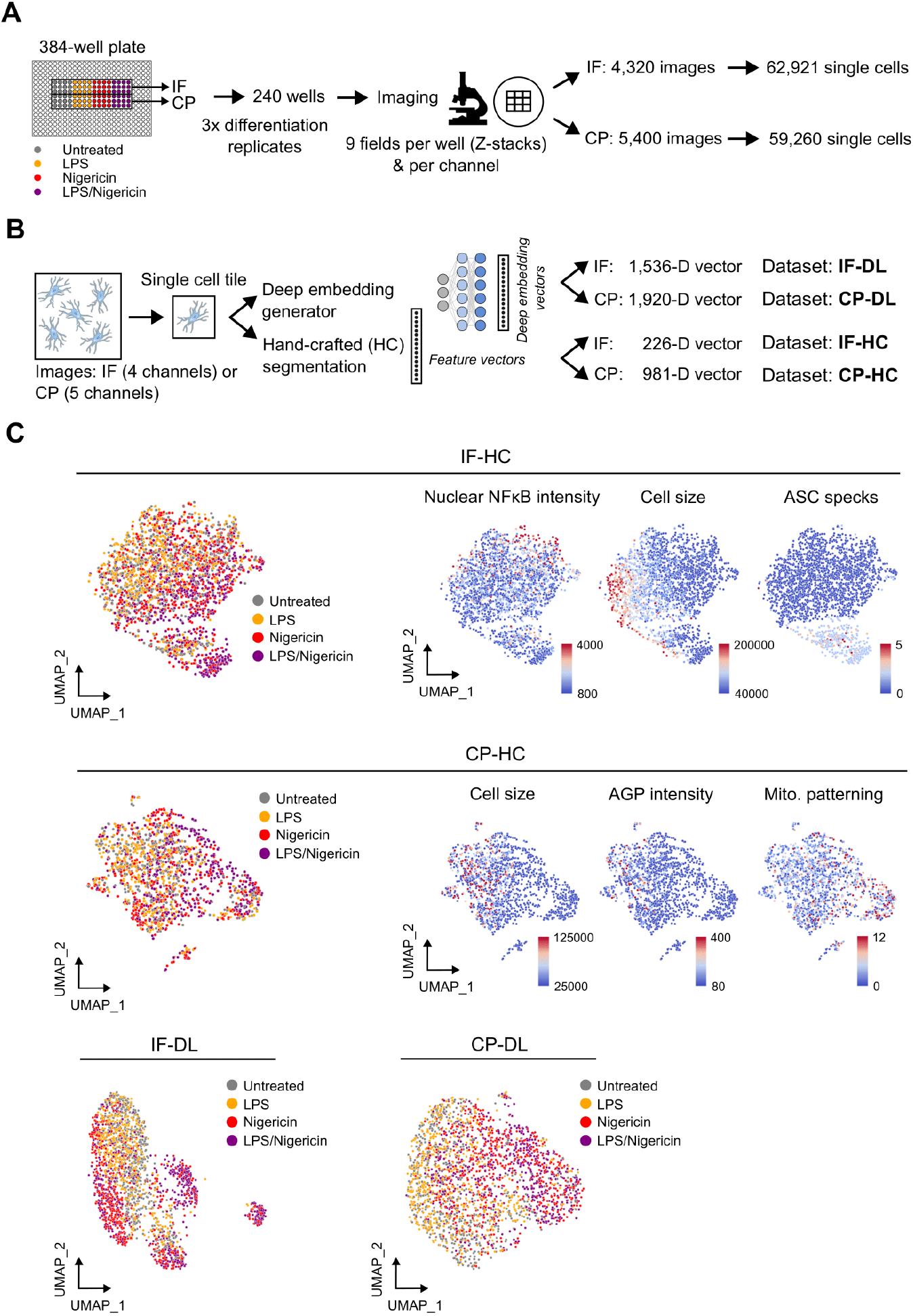
Experimental workflow and visualization of single-cell phenotypic landscapes. **(A)** Overview of the experimental setup. iPSC-derived microglia were seeded in 384-well plates and treated with four conditions (Untreated, LPS, Nigericin, LPS/Nigericin). Cells were stained using either immunofluorescence (IF) or Cell Painting (CP) protocols and imaged across 240 wells, yielding over 62,921 single cells for IF and 59,260 for CP. **(B)** Image analysis pipeline for generating single-cell feature vectors. Each segmented cell tile was processed using two parallel approaches: handcrafted feature extraction and deep learning-based embedding generation. This resulted in four datasets: IF_HC (226 features), CP_HC (981 features), IF_DL (1,536-dimensional embeddings), and CP_DL (1,920-dimensional embeddings). **(C)** UMAP projections of the four datasets, colored by treatment condition. Distinct phenotypic clusters corresponding to activation states were observed across all datasets.

Mean per-image values often obscure microglial heterogeneity and dynamic subpopulation shifts in response to inflammatory stimuli. To resolve this, we extracted single-cell quantitative data using IF and CP, capturing both targeted molecular markers and broad morphological features. For both datasets, single-nucleus segmentation was performed to isolate individual cells, enabling downstream extraction of quantitative features from each cell tile (**Figure 2B**).

Two distinct image analysis methods were applied to extract quantitative features from segmented or unsegmented single-cell images. The first analysis approach involved handcrafted feature extraction, which relies on image thresholding and the segmentation and quantification of experimenter-determined structures. Handcrafted features quantify interpretable cellular attributes, such as shape, size, intensity, and texture. For IF images, 226 features were extracted which were specifically designed to capture features linked to the used fluorescent stains Hoechst, NF-κB, CD45 and ASC. These for example included the area and intensity of CD45 positive cells and ASC specks or the intensity of NF-κB in different cellular compartments. CP images yielded 981 features, reflecting the broader range of compartments labeled by this technique. CP-derived features do not quantify specific aspects microglia biology but rather measure various aspects of general cell biology linked to the nucleus, cytoplasm, mitochondria, endoplasmic reticulum (ER), Golgi apparatus, and the actin cytoskeleton (**Table S1, Figure 2B**).

The second analysis approach did not rely on image segmentation but utilized deep learning-based feature extraction via the self-supervised vision transformer model DINOv2 (Self-Distillation with No Labels) ^23^. Unlike segmentation- and thresholding-based methods, DINOv2 learns high-dimensional representations of cellular morphology directly from image data without requiring labeled training sets. It captures complex, non-linear patterns in cellular organization and texture that may be too subtle for traditional algorithms. This method produced 384 features per imaging channel, resulting in 1,536-dimensional embeddings for IF images (4×384) and 1,920 for CP images (5x384), enabling rich, unbiased phenotypic profiling (**Figure 2B**).

To visualize the high-dimensional phenotypic landscape of microglial activation, we applied Uniform Manifold Approximation and Projection (UMAP) to the four generated single-cell feature sets: immunofluorescence with handcrafted features (IF_HC), immunofluorescence with deep learning embeddings (IF_DL), Cell Painting with handcrafted features (CP_HC), and Cell Painting with deep learning embeddings (CP_DL) (**Figure 2C**). Overlaying treatment conditions onto the UMAP projections revealed well-separated iMG subpopulations that corresponded to distinct activation states, consistently across all four feature extraction approaches (**Figure 2C**). This reproducibility highlights the robustness of both the staining modalities, targeted IF and unbiased CP, and the computational pipelines, including handcrafted and deep learning-based features, in capturing biologically meaningful phenotypic diversity. The convergence of these orthogonal methods underscores the reliability of the observed activation landscape and supports the use of this framework for high-resolution microglial state profiling.

Additionally, the same selected handcrafted feature values as in **Figure 1D-E** were overlaid on the respective UMAP projections. For example, CD45 staining-based cell surface was lowest in iMG originating from Nigericin and LPS/Nigericin treated wells. Similarly, the number of NLRP3 inflammasome activation-related ASC specks was highest in fully primed and activated cells, although also some iMG from untreated wells showed activation-related phenotypes (**Figure 2C**). CP-derived features showed a similar, but less pronounced correlation with iMG activation treatment (**Figure 2C**).

### Phenotypic clustering and classification of iMG activation states

To robustly cluster high-dimensional single-cell phenotypic data, we employed two complementary algorithms: the graph-based Leiden algorithm, widely used in scRNA-seq analysis, and the Gaussian Mixture Model (GMM), a probabilistic soft clustering method, that unlike graph-based methods, is designed to model the probability of a cell belonging to a cluster. Despite their proven utility in various biological data contexts both algorithms are rarely applied to single-cell high-content imaging data.

To investigate which algorithm is better suited for high-dimensional single-cell phenotypic data we designed a benchmarking pipeline (**Figure 3A**). Starting with image segmentation and deep learning-based feature extraction, the data underwent Z-score normalization, feature filtering, and principal component analysis (PCA). The two clustering algorithms, GMM and Leiden, were then applied independently to generate cluster assignments, which were aggregated per well into Cluster Composition Vectors (CCVs). These CCVs, representing the proportion of cells per cluster, served as input for supervised classifiers trained to predict treatment conditions. Classifier performance was evaluated using F1 scores based on the known used iMG treatments, enabling a direct comparison of clustering accuracy and reproducibility between GMM and Leiden across multiple datasets and conditions (**Figure 3B**). This approach allowed us to quantify how well each clustering method captured biologically meaningful phenotypic variation.

**Figure 3.**
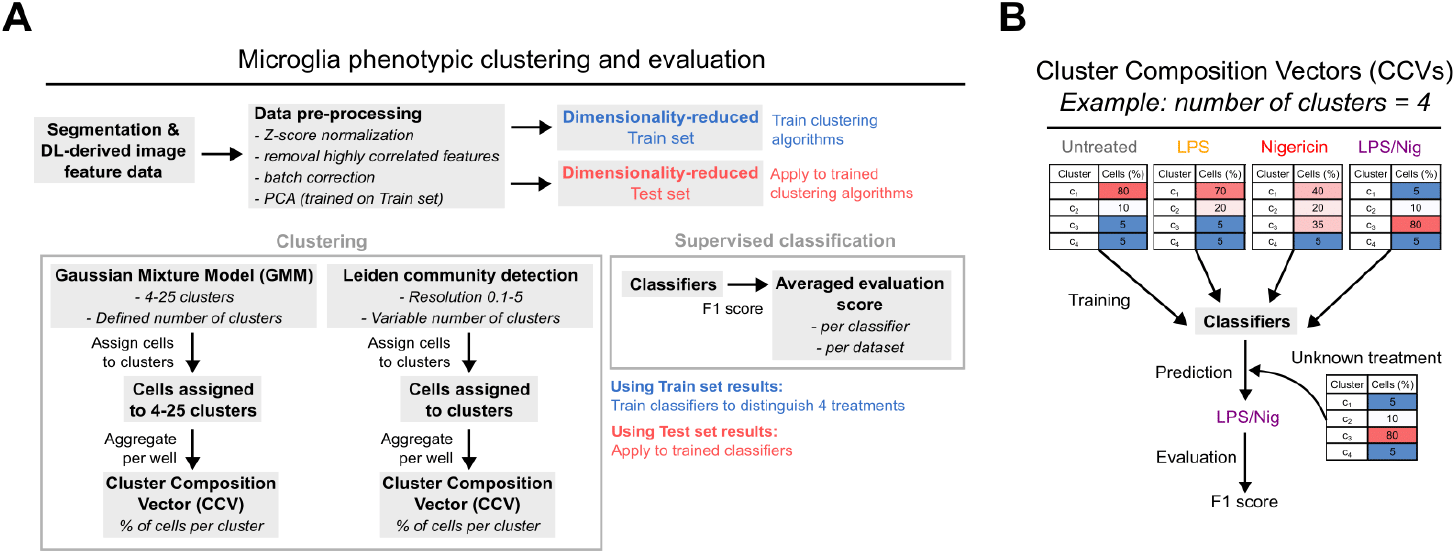
Workflow for microglial phenotypic clustering and classification using imaging-derived features. **(A)** Schematic overview of the clustering and evaluation pipeline. Deep learning-derived single-cell image features underwent preprocessing steps including Z-score normalization, removal of highly correlated features, and PCA. Clustering was performed using either Gaussian Mixture Models (GMMs) with a defined number of clusters ranging from 4-25 or Leiden community detection with a variable resolution parameter ranging from 0.1-5. Cluster assignments were aggregated per well into Cluster Composition Vectors (CCVs), representing the proportion of cells per cluster. These CCVs were used to train supervised classifiers on the training set and evaluate performance on the test set using F1 scores. **(B)** Example of CCVs for four clusters across different treatment conditions (Untreated, LPS, Nigericin, LPS/Nigericin). The relative abundance of clusters varies by treatment, forming the basis for supervised classification and enabling quantitative assessment of clustering performance.

### Deep learning-based feature extraction is suitable for detection of iMG activation states

To assess the robustness of our phenotyping framework, we evaluated how well CCVs derived from GMM and Leiden clustering could be used to predict treatment conditions using supervised classifiers. CCVs effectively serve as a “phenotypic fingerprint” for each cell. Across all datasets, deep learning-derived features consistently outperformed handcrafted features in classification accuracy, with F1 scores exceeding 0.8 for IF_DL and CP_DL datasets (**Figure 4A**). In contrast, handcrafted features yielded lower predictive performance, particularly for CP_HC, which showed the lowest classification accuracy. The superior performance of DL data likely stems from the ability of ViT models to capture subtle, non-linear morphological patterns that are not easily encoded by predefined feature sets.

**Figure 4.**
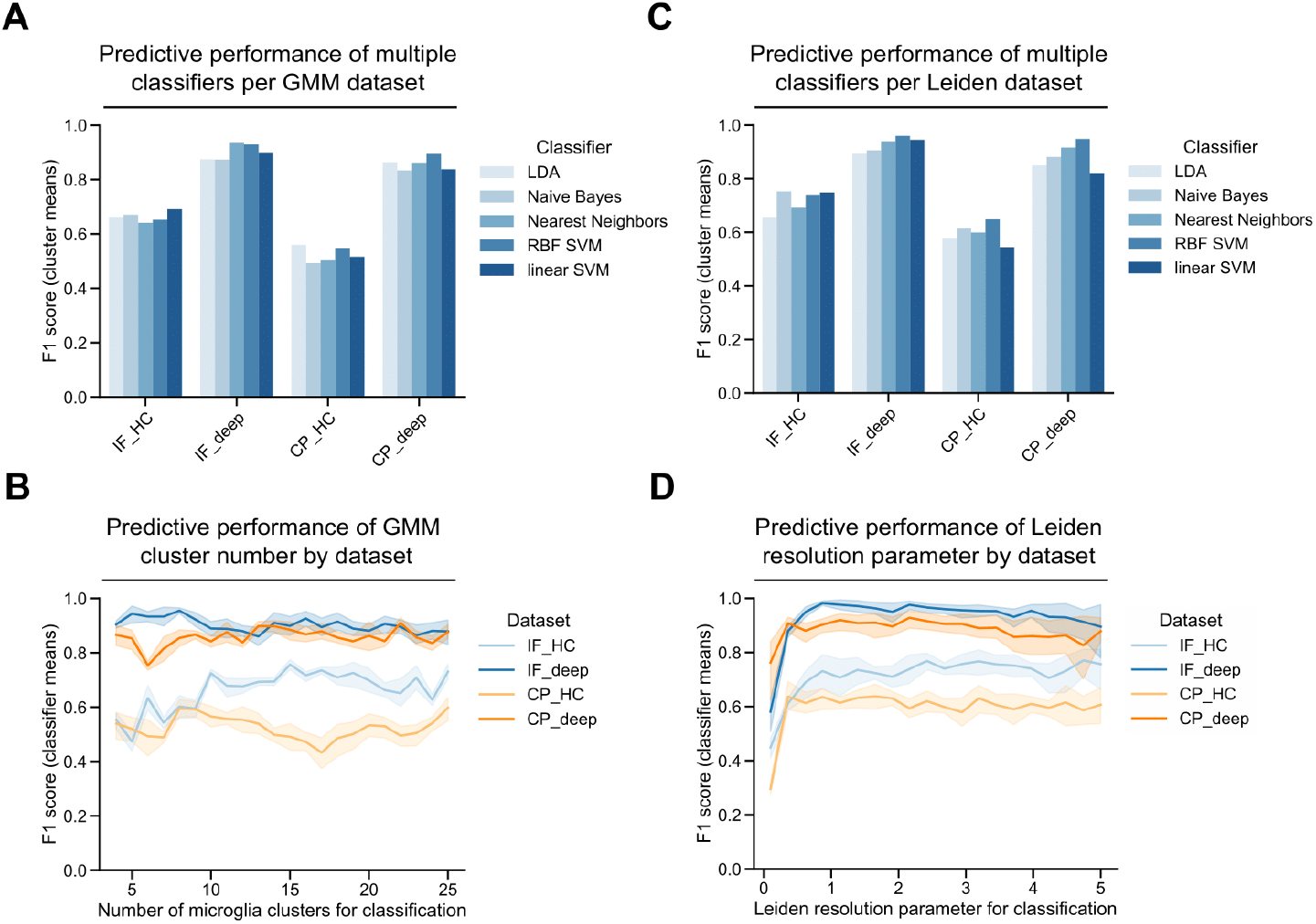
Classifier performance across clustering methods, feature types, and parameter settings. **(A)** Supervised classification performance (F1 scores) using Cluster Composition Vectors (CCVs) derived from GMM clustering across four datasets: immunofluorescence with handcrafted features (IF_HC), immunofluorescence with deep learning features (IF_deep), Cell Painting with handcrafted features (CP_HC), and Cell Painting with deep learning features (CP_deep). **(B)** Effect of GMM cluster number variation on classification accuracy. **(C)** Classifier performance using CCVs derived from Leiden clustering. **(D)** Effect of the Leiden resolution parameter on classification accuracy.

We next examined how the number of GMM clusters influenced classifier performance. Using low cluster numbers (e.g., 4–6) led to reduced F1 scores for IF-HC features, while varying the cluster number caused highly variable performance for CP-HC features, indicating that these configurations failed to capture the full morphological diversity of microglial states (**Figure 4B**). Deep learning-derived features were more resilient to variations in cluster number, maintaining high predictive accuracy even at lower resolutions.

To benchmark against graph-based clustering, we repeated the classification analysis using CCVs derived from Leiden clustering. The overall trends mirrored those observed with GMM: deep learning features again outperformed handcrafted ones, and Naïve Bayes and Nearest Neighbors classifiers consistently underperformed relative to linear and RBF SVMs (**Figure 4C**). Finally, we evaluated the effect of the Leiden resolution parameter on classification performance. Similar to GMM cluster number, low resolution values led to diminished F1 scores, particularly for handcrafted features, while deep learning-derived features remained robust across a wide range of resolution settings (**Figure 4D**).

### The optimal GMM cluster number depends on the feature data origin

To determine the optimal number of clusters for each dataset, we applied GMMs with varying cluster counts and evaluated model fit using the Akaike Information Criterion (AIC). The AIC curves revealed dataset-specific optima, with the lowest AIC values corresponding to 16 clusters for the IF_HC dataset, 15 clusters for IF_DL, 19 clusters for CP_HC, and 16 for CP_DL (**Figure 5A**). These values were used as the final cluster numbers for downstream analyses.

**Figure 5.**
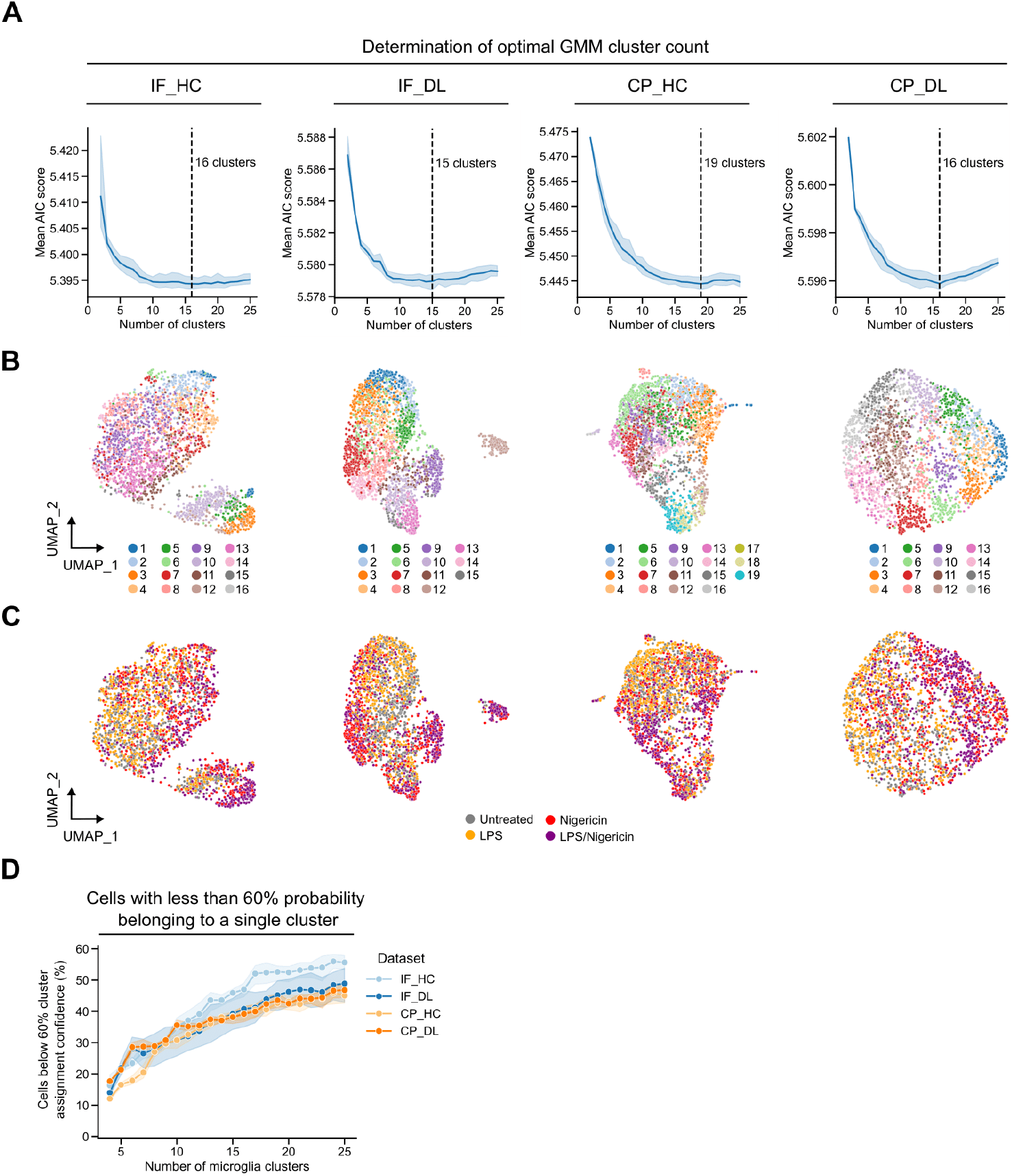
Determination of optimal GMM cluster number. **(A)** Model selection using Akaike Information Criterion (AIC) to determine the optimal number of GMM clusters for each dataset. The lowest AIC scores indicated optimal cluster counts of 16 for IF_HC, 15 for IF_DL, 19 for CP_HC, and 16 for CP_DL. **(B)** UMAP projections of single-cell data colored by GMM cluster assignments for each dataset. **(C)** UMAP projections of the same datasets colored by treatment condition (Untreated, LPS, Nigericin, LPS/Nigericin). **(D)** Proportion of cells with ambiguous cluster assignments (less than 60% probability of belonging to a single cluster) plotted against the number of clusters for each dataset.

To visualize the phenotypic landscape captured by GMM clustering, we projected the single-cell data into two dimensions using UMAP and colored cells by their assigned cluster labels. This revealed distinct and well-separated phenotypic subpopulations across all four datasets, with deep learning-derived features (IF_DL and CP_DL) showing more compact and structured cluster distributions compared to their handcrafted counterparts (**Figure 5B**). These results suggest that deep learning embeddings capture more coherent and biologically meaningful phenotypic variation.

We next examined how treatment conditions were distributed across the phenotypic space. UMAP projections colored by treatment revealed that LPS/Nigericin-treated cells occupied distinct regions in all datasets, particularly in IF_DL and IF_HC, where activated phenotypes formed clearly separated clusters (**Figure 5C**). Nigericin and LPS alone also induced partial shifts in the phenotypic landscape, while untreated cells were more broadly distributed. These findings show that our imaging-based phenotyping framework can resolve treatment-induced microglial heterogeneity.

Interestingly, we observed that increasing the number of clusters using GMM led to a higher proportion of microglia cells that could not be confidently assigned to any cluster at a 60% probability threshold (**Figure 5D**). This counterintuitive result arises from the probabilistic nature of GMMs, which assign each cell a set of probabilities corresponding to its likelihood of belonging to each cluster. As the number of clusters increases, each cluster becomes more narrowly defined, reducing the maximum probability that a given cell belongs to any one cluster. Additionally, the total probability mass is distributed across more components, often resulting in lower individual assignment probabilities. Consequently, more cells fall below the confidence threshold and remain unassigned.

### Morphology-based iMG cluster composition changes across iMG activation states

Since the predictive performance was highest for the IF_DL dataset, we examined in detail how the 15 identified clusters varied across different iMG priming and activation conditions. Cluster prevalence shifted markedly between treatments: untreated iMG were broadly distributed, while LPS and Nigericin exposure led to enrichment in clusters 1, 3, and 10, and clusters 7 and 14, respectively. The combined LPS/Nigericin treatment induced the most pronounced redistribution, with clusters 7, 9, and 12 becoming highly enriched and others nearly absent (**Figure 6A**). These changes indicate that specific morphological phenotypes correspond to distinct activation states, and that GMM clustering sensitively captures these transitions.

**Figure 6.**
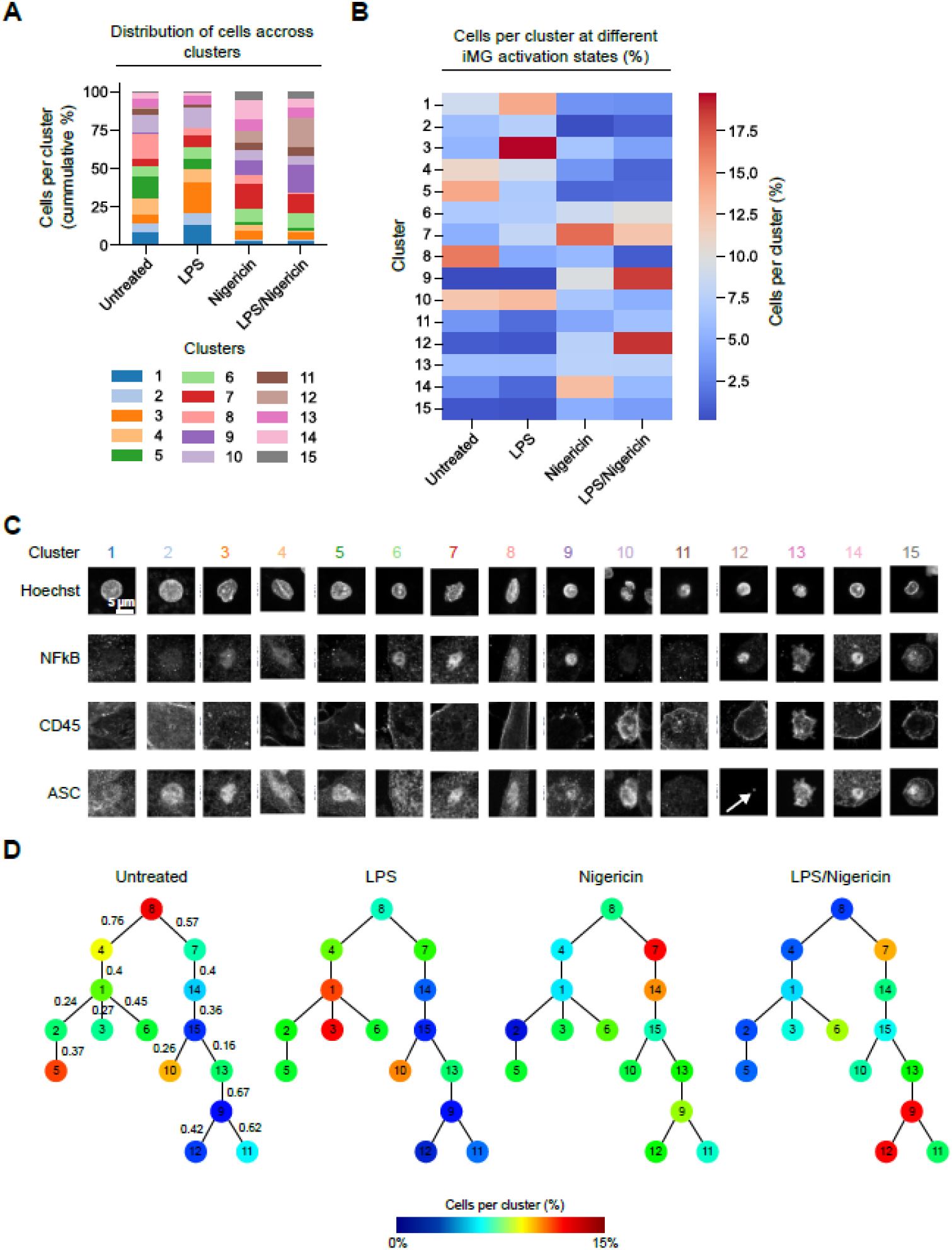
Morphology-based iMG cluster composition changes across iMG activation states based on IF_DL data. **(A)** Bar graph showing the distribution of cells across clusters under four treatment conditions: Untreated, LPS, Nigericin, and LPS/Nigericin. Clusters are color-coded as indicated in the legend. **(B)** Heatmap displaying the percentage of cells per cluster across the same four conditions, with a color gradient from blue to red representing increasing percentages. **(C)** Representative images of cells from clusters 1 to 15, stained with Hoechst, NFkB, CD45, and ASC. **(D)** Minimum spanning tree (MST)-based graphs to illustrate cluster relationships. Cosine distances were used as edge weights and are shown only in the left-most graph for clarity, as they are identical across all graphs. Graph nodes are color-coded to reflect the percentage of cells within each cluster, ranging from 0% (blue) to 15% (red).

To visualize cluster redistribution across conditions, we plotted cluster frequencies as a heatmap (**Figure 6B**). This representation highlights condition-specific enrichment patterns: for instance, cluster 3 is almost exclusively present in LPS-primed iMG, while clusters 2, 6, 11, 13, and 15 remain relatively stable across treatments. These stable clusters may represent non-responsive or dying cells. To assess clustering confidence, we visualized only cells with a high assignment probability (>90%) (**Figure 6C**). Clusters 5 and 8, enriched in untreated iMG, exhibit large nuclei and cell bodies, diffuse NFκB localization, and absence of ASC specks—features consistent with a non-activated state. In contrast, cluster 12, enriched in LPS/Nigericin-treated cells, shows small cell size, nuclear NF-κB localization, and prominent ASC specks, indicative of full activation. Cluster 9 shares features with cluster 12 but lacks ASC specks, suggesting an intermediate activation state. Cluster 13, stable across all conditions and marked by fragmented nuclei and cytoplasm, likely represents dead cells.

To explore phenotypic relationships and transitions, we constructed minimum spanning trees (MSTs) using cluster centroids in PCA space, with cosine distances as edge weights (**Figure 6D**). MSTs reveal efficient, cycle-free connections between phenotypically similar clusters. Separate MSTs for each condition show treatment-specific structural patterns: clusters 1–6 form a distinct branch associated with untreated or LPS-primed iMG, while clusters 9–15 form a separate branch enriched in Nigericin-activated or fully LPS/Nigericin-primed cells. Cluster 13 appears consistently across conditions, reinforcing its potential link to cell death. The presented approach to measure and classify iMG phenotypes might guide the identification of bioactive molecules that selectively influence desired iMG phenotypes, such as those associated with early activation stages (clusters 1–6) or later, fully activated states (clusters 9–15).

## Discussion

Our study establishes a probabilistic imaging framework that captures the continuous and heterogeneous nature of microglial activation. By applying GMMs to high-content imaging data, we move beyond discrete classifications to model the probability of a cell existing in a given state, thereby revealing transitional populations that are likely critical for understanding neuroinflammatory dynamics. This approach to deconstruct microglial heterogeneity complements our and others previous work using conditional generative models to reveal invisible cell phenotypes, together contributing to a growing set of AI-based tools for identifying and characterizing novel cellular states in health and disease ^24–26^. Unlike previous morphology-driven microglia classification approaches that rely on low-magnification brain slice imaging ^27–29^ or non-scalable modalities such as fluorescence lifetime imaging ^30^, our study aimed to detect microglial subpopulations using high-content, high-throughput compatible 2D cell culture assays with physiologically relevant iMG (**Figure 1**). We developed a robust single-cell image analysis pipeline that ensures stable feature extraction using either a deep learning or a handcrafted approach across multiple iMG differentiation batches and is compatible with various staining approaches such as IF and Cell Painting (**Figure 2A-B**). We benchmarked GMMs against the commonly used Leiden algorithm to allow the detection of both frequent and rare microglial phenotypes (**Figure 3**).

Clustering biological data or detecting communities within it is a challenging task, primarily because ground truth labels are often unavailable. Additionally, the granularity of clusters can be chosen arbitrarily: data may be grouped into a few broad categories or divided into many fine-grained subgroups ^13,31^. Among the most widely used clustering approaches in single-cell analysis are the Louvain and Leiden algorithms ^21,32,33^. These methods operate on a graph-based representation of the data, typically constructed using a k-nearest neighbor (kNN) graph. Louvain clustering optimizes modularity to detect communities, while Leiden improves upon this by refining partition quality and ensuring well-connected clusters ^21^. Despite their popularity in transcriptomic analyses, these algorithms are less suitable for phenotypic imaging-derived feature data. This is primarily because they rely on graph topology rather than the actual distribution of features in Euclidean space, making them sensitive to graph construction parameters and less capable of capturing the continuous and overlapping nature of morphological phenotypes.

Alternative clustering methods such as K-means and GMMs offer more appropriate frameworks for analyzing imaging-derived features ^13^. K-means partitions data into a predefined number of clusters by minimizing within-cluster variance, assuming spherical clusters of similar size. However, it lacks flexibility in modeling clusters with varying shapes and densities, and rare populations can be easily overlooked ^31^. In contrast, GMMs model the data as a mixture of multivariate Gaussian distributions, allowing for elliptical clusters with different orientations and covariances. Importantly, GMMs provide probabilistic cluster assignments, which are particularly valuable for capturing transitional states in biological systems. We decided to test GMM-based clustering since it has been applied before to define “communities” based on image-derived feature data. GMM clustering has been used to identify bacterial subpopulations, classify particles after cryogenic electron microscopy (cryo-EM), or seed image classification in plant breeding applications ^34–36^. Additionally, GMM clustering has been applied to human cell-derived imaging data to characterize subtle phenotypic changes upon drug treatment ^37,38^ or the classification of morphological heterogeneous cells based on cell shape ^39–41^.

However, GMM-based clustering also has limitations: it assumes that the data can be well-approximated by a mixture of Gaussian distributions, which may not hold for complex or non-Gaussian feature spaces. Additionally, it can be sensitive to initialization and the choice of the number of components and may struggle with high-dimensional data where covariance estimation becomes unstable. We addressed these limitations by implementing several strategies to enhance the robustness and interpretability of GMM clustering in the context of high-dimensional imaging-derived data (**Figure 3**). First, we applied dimensionality reduction via PCA prior to clustering, which mitigates the curse of dimensionality and stabilizes covariance estimation by reducing noise and redundancy in the feature space. Second, we used the Akaike Information Criterion (AIC) to systematically determine the optimal number of clusters for each dataset, avoiding arbitrary selection and overfitting (**Figure 5A**). Third, we introduced a probabilistic confidence threshold (e.g., 60%) to filter out ambiguous cluster assignments, thereby focusing downstream analyses on high-confidence phenotypic states. This approach allowed us to retain the interpretability and flexibility of soft clustering while minimizing the impact of uncertain assignments. Despite these mitigations, as the number of clusters increases, the model tends to assign lower maximum probabilities to each cell, leading to a higher proportion of unassigned cells (**Figure 5D**). This phenomenon reflects the trade-off between resolution and confidence in soft clustering and underscores the importance of integrating GMM outputs with biological validation and orthogonal readouts.

The performance of clustering and classification was strongly influenced by the choice of feature extraction method and staining modality. Across all datasets, deep learning-derived features consistently outperformed handcrafted features in predicting microglial activation states (**Figure 4**). This likely reflects the ability of self-supervised models like DINOv2 to capture subtle, non-linear morphological patterns that are difficult to encode using predefined metrics. These embeddings likely integrate complex spatial relationships and textural cues that are biologically meaningful but not easily quantifiable through classical image analysis. Among the staining modalities, immunofluorescence features, particularly those derived from deep learning, yielded the highest classification accuracy. This can be attributed to the targeted nature of immunofluorescence, which includes mechanistically relevant markers such as NF-κB nuclear translocation and ASC speck formation, both of which are directly linked to NLRP3 inflammasome activation. These markers provide strong, interpretable signals that correlate with functional microglial states, enhancing the biological relevance of the extracted features (**Figure 6**). In contrast, Cell Painting, while offering broader morphological coverage, lacks microglia-specific markers and may dilute condition-specific signals with unrelated structural variation.

Microglia-driven neuroinflammation is central to many neurological disorders, including Alzheimer’s, Parkinson’s, multiple sclerosis, and ALS. Single-cell RNA sequencing has revealed distinct microglial subtypes, such as DAMs, potentially offering new therapeutic targets ^5,9^. iMG, while not identical to primary human microglia, closely mimic their transcriptional and functional profiles and can be produced at scale, making them ideal for high-throughput drug screening, unlike limited primary microglia ^14,42,43^. Importantly, Sun *et al*. recently showed that iMG showed similar microglial state dynamics than primary microglia from human donors ^44^.

While the primary objective of this study was to establish a technical framework for microglia characterization, one limitation is the reliance on LPS and Nigericin for microglial stimulation, which may not fully capture the diversity of microglial activation states. In the future it will be interesting to determine if stimuli linked to brain challenges such as amyloid or α-synuclein deposition, infected, damaged, or degenerating neurons will lead to a different community structure ^2^. Furthermore, it will be crucial to understand how variable phenotypic responses are within iMG from various healthy donors, but also versus iMG from donors carrying mutations linked to neurodegenerative or neurodevelopmental diseases.

Despite the various and highly dynamic contexts in which microglia exist, our findings suggest that combining targeted IF staining with DL-based feature extraction provides a sensitive and specific approach for resolving microglial activation states. This synergy enables the detection of both subtle and pronounced phenotypic shifts, supporting the use of similar pipelines in future high-throughput screening applications aimed at identifying modulators of microglial function.

## Materials and Methods

### Microglia differentiation

iPSCs were differentiated by LIFE & BRAIN into iMG as described previously ^14^ (patent: WO2021180781A ^45^). In brief, embryoid bodies (EBs) were cultured in suspension on an agitator in EB basal media containing N2 and B27 supplements (1X; Gibco) with stepwise supplementation of BMP4, FGF2, Activin A, and a WNT inhibitor. On day 4, EBs were transferred onto poly-L-ornithine/fibronectin-coated nylon mesh macrocarriers to promote progenitor outgrowth, followed by suspension culture in the presence of IL-3, IL-34, and M-CSF.

### Media preparation

The seeding medium consisted of DMEM-F12 with HEPES (ThermoFisher), supplemented with GlutaMAX (1X), N2 supplement (1X), and freshly added recombinant human IL-34 (100 ng/mL). For priming and activation steps, IL-34 was omitted

### Plate coating

384-well plates (Perkin Elmer) were coated with Poly-L-Lysine (PLL) hydrobromide (Sigma/Merck) at 5 µg/mL (1:200 dilution in sterile distilled water). Plates were incubated overnight at 37 °C with 5% CO_2_. PLL coating dispense was performed with a multichannel pipet. Before cell seeding, the coating was washed twice with 50µL sterile PBS/well using a BioTek EL406 Washer Dispenser (Agilent). 10µL PBS remained in each well after the final wash.

### Cell seeding and treatment

Cells harvested from the supernatant were centrifuged (400g, 5 min), filtered through a 40 μm cell strainer and counted using the Countess Automated Cell Counter (Invitrogen) with Trypan Blue exclusion. A suspension of 160,000 cells/mL was prepared, and 50 µL (8,000 cells) was seeded per well in DMEM-F12 with HEPES (ThermoFisher), supplemented with GlutaMAX (1X), N2 supplement (1X), and freshly added recombinant human IL-34 (100 ng/mL). Cells were primed with 1 µg/mL LPS (Invitrogen) in IL-34 free medium. After 1 hour of LPS priming, activation was performed by adding 10 µL of 70 µM Nigericin (final concentration 10 µM) and incubation for an additional 3 hours. Both priming and activation steps were performed by using a Vprep pipetting system (Agilent).

### Fixation and immunostaining

After priming and activation, cells were fixed with 4% paraformaldehyde (Euromedex) for 30 minutes at room temperature and washed with PBS using BioTek EL406 Washer Dispenser (Agilent). Blocking and permeabilization were performed using a solution of 0.3% Triton X-100 and 10% FBS in PBS. Primary antibodies (CD45, ASC, NFκB p65) were incubated overnight at 4 °C. After washing, secondary antibodies conjugated with Alexa Fluor dyes (AF488, AF594, AF647) and Hoechst 33342 were applied for 2 hours at room temperature in the dark. Plates were sealed and stored at 4 °C until imaging.

### Cell Painting

After priming and activation, cells were fixed with 4% paraformaldehyde (Euromedex) for 30 minutes at room temperature and washed with PBS using BioTek EL406 Washer Dispenser (Agilent). Prior to staining, 10 µL of medium was removed from each well, leaving 10 µL. The PhenoVue™ Cell Painting kit (Revity) was used. A staining solution was prepared at 1.5x concentration in a total volume of 3 mL using the following components: PhenoVue 555 WGA (2.25 µg/mL), PhenoVue Fluor 488 Concanavalin A (7.5 µg/mL), PhenoVue Fluor 568 Phalloidin (12.375 nM), PhenoVue Hoechst 33342 Nuclear Stain (1.5 µg/mL), PhenoVue 512 Nucleic Acid Stain (9 µM), PhenoVue Dye Diluent A (1.5x), and dH_2_O. Each well received 20 µL of the 1.5x staining solution. Plates were incubated in the dark at room temperature for 30 minutes. Following incubation, wells were manually washed three times with PBS. Plates were sealed with aluminum foil and stored at 4 °C until imaging.

### Imaging

Imaging was performed on a Cell Voyager 7000 system (Yokogawa) microscope in scanning confocal mode using a dual Nipkow disk and a cooled sCMOS camera with 2,560×2,160 pixels and a pixel size of 6.5 μm without pixel binning. The system’s CellVoyager software, the 405/488/561/640-nm solid laser lines, and a dry 40X objective were used to acquire 9 images in a 3×3 orientation from three 1 µm-separated Z-planes from the center of each well. Wells were visited in a row-wise “zigzag” fashion. Maximum intensity projections were performed.

### Feature extraction

#### DINOv2 feature extraction

The feature extraction procedure consists of two main stages: nuclei segmentation and cell cropping, followed by feature extraction using a pretrained vision transformer model.

In the first stage, nuclei were segmented based on intensity, size, and density thresholds. Once the nuclei were identified, square image patches of size 140×140 pixels were cropped around each nucleus, with the nucleus center positioned at the center of the crop. The crop size was determined through visual inspection to ensure sufficient coverage of the cellular region surrounding each nucleus. For feature extraction, we employed DINOv2, a self-supervised Vision Transformer (ViT) model trained on millions of images to generate general-purpose visual representations for downstream tasks ^23^. The model was converted to ONNX Runtime and integrated into our image analysis platform Phenolink, enabling efficient large-scale feature extraction. Each cropped single-channel image was triplicated to create pseudo RGB channels to form a 3×140×140pixel input compatible with the DINOv2 model. The model outputs a 384-dimensional feature vector per input crop. Accordingly, we obtained 1536 features per cell for immunofluorescence (IF) staining (4 channels) and 1920 features per cell for Cell Painting staining (5 channels).

#### PhenoLink feature extraction

For IF data, image segmentation and feature extraction was performed with our in-house software PhenoLink (https://github.com/Ksilink/PhenoLink). Image segmentation was performed on single cell-tiled images. Using intensity and size thresholds specific to each of the four imaging channels (DNA, NF-κB, CD45, and ASC) we extracted a total of 226 morphological features, including count, texture, context, intensity and shape features (Table S1). We used location constraints to identify subcellular compartments. For example, by subtracting the segmented nucleus from the whole cell we obtained the cytoplasm mask.

#### Cell Profiler feature extraction

We processed Cell Painting images using a CellProfiler pipeline adapted from the JUMP-Consortium standard workflow ^46^. Nuclei were segmented as primary objects, and non-round or dead nuclei were excluded based on shape and intensity thresholds. The pipeline then identified cell and cytoplasm boundaries, enabling extraction of per-cell morphological features. These included measurements of fluorescence intensity, texture, granularity, shape, and area across all imaging channels (Table S1). Following feature extraction, single-cell profiles were aggregated per well, normalized, and outliers were removed to ensure robust data quality for downstream analysis.

### Data pre-processing

The above steps were applied consistently across all datasets. Wells identified as corrupt or empty during image acquisition were excluded from further analysis. To account for potential batch effects and ensure comparability across features, we performed per-plate normalization, transforming each feature to have zero mean and unit standard deviation. This step helped to reduce mild plate effects and brought all features into the same numerical range. One of the experiments included approximately three times more wells than the others. To mitigate the resulting data imbalance, we randomly subsampled the other two experiments, yielding a more balanced number of samples across experiments. For datasets involving handcrafted features (e.g., extracted via CellProfiler or Phenolink), we further removed highly correlated features (more than 90 percent Pearson correlated features) to reduce redundancy and potential noise in downstream analyses.

### Data clustering and classification

To assess the performance of each dataset in a multi-class classification task involving four experimental conditions (Unactivated, LPS, Nigericin, and LPS/Nigericin), we designed a cross-validated classification pipeline. For each dataset, one of the three experiments was designated as the test set, while the remaining two were used for training. This process was repeated three times, rotating the test experiment each time. The final performance metrics were averaged across the three folds, allowing for a robust comparison across datasets and simulating generalization to unseen data. This setup helps evaluate whether a dataset contains unbiased and generalizable information for downstream classification tasks. Following the train-test split, the training data was first used to fit a Principal Component Analysis (PCA) model, reducing the dimensionality to 10 components. Next, clustering was performed on the PCA-reduced data to identify putative cell types using two distinct methods: Gaussian Mixture Models (GMM) and Leiden community detection. Since the true number of cell types was unknown, we explored a range of cluster numbers by fitting 22 GMM models with cluster counts ranging from 4 to 25. To ensure confidence in cluster assignments, we applied a probability threshold of 0.7, below which cells were labeled as belonging to an unknown cluster (cluster ID: –1). For the Leiden method, we varied the resolution parameter from 0.1 to 5.0, resulting in 20 different clustering models. For Leiden clustering in the test set, cluster labels were assigned using a k-nearest neighbors (KNN) approach with *k* = 15 and cosine distance as the similarity metric. For each well, we then computed a Cluster Composition Vector (CCV), a vector representing the proportion of cells within that well assigned to each cluster. These CCVs served as input features for a classifier trained to distinguish the four control conditions. As classifiers, we employed five different algorithms: Support Vector Machine (SVM) with L2 penalty, K-nearest neighbors (KNN) with *k* = 3, SVM with an RBF kernel, Linear Discriminant Analysis (LDA), and Naive Bayes. The trained PCA, clustering model, and classifiers were then applied to the test set. The F1 score of the classification on the test set was used as the evaluation metric.

### Activation state detection

The objective of this analysis was to uncover and characterize the diversity of cell types present in our dataset. To maximize the amount of information available, we merged all experimental data prior to splitting into training and testing sets, unlike in a supervised classification setting. The dataset was then split into 70% training and 30% testing data. Although the pipeline follows a structure similar to that used in classification tasks, several key differences apply here: To determine the optimal number of distinct cell types, we applied Gaussian Mixture Models (GMMs) with the number of clusters ranging from 2 to 25. For each model, we computed the Akaike Information Criterion (AIC), selecting the number of clusters that minimized the AIC score as the optimal solution. For the visualization of single iMG per cluster, we retained only the cells assigned to a specific cluster with a posterior probability of >0.9, ensuring high-confidence cluster memberships. To visualize the trajectory of cell states, from untreated to LPS/Nigericin activated phenotypes, we constructed a graph based on a minimum spanning tree (MST). Nodes in this graph correspond to the cluster centroids in PCA space, and edge weights were defined as the cosine distance between cluster centers. The resulting tree was rearranged to position the most inactivated cluster at the top. We defined the most inactivated cluster as the one with the largest positive difference between the proportion of its cells originating from untreated wells versus LPS/Nigericin-treated wells. This allowed us to infer a progression of cell states along the graph structure.

### Statistics and data plotting

All statistical analyses and visualizations were performed in Python using Seaborn and Matplotlib with default statistical settings. SuperPlots, a variation of boxplots, were used to display well-level averages (mean of 9 images per well) showing median, interquartile range, and whiskers at 1.5×IQR, with the mean value of each biological replicate (independent iMG differentiation) overlaid as larger markers ^47^. Bar graphs and line plots show means, while line plots include also shaded 95% confidence intervals. UMAP scatter plots were used to visualize single-cell data and were randomly down sampled to 20% to improve clarity; each point represents an individual cell, whereas all other plots are based on well-level averages. Data from three independent iMG differentiation batches were used throughout, and statistical comparisons were performed using two-sided independent t-tests on biological replicate means using the Python “statannotations” package ^48^.

## Data and code availability

All raw data and computational scripts used to perform the analyses presented in this study, including Python Jupyter notebooks, are publicly available on GitHub at https://github.com/Ksilink/microglia-morphological-profiling.

## Acknowledgements

The authors thank all colleagues at Ksilink and LIFE & BRAIN GmbH who contributed through continued discussions and feedback to the advancement of the project. This project was jointly funded by Zentrales Innovationsprogramm Mittelstand (ZIM) of the German Federal Ministry for Economic Affairs and Energy (KK5450601SK2 to LIFE & BRAIN) and by the Banque publique d’investissement (Bpifrance) (DOS0193862/00 to Ksilink).

## Author contributions

KA designed, implemented, and performed image and data analysis, AW designed and performed experiments, MM and BB developed the iMG priming and activation scheme, MS performed iMG differentiation, ZH performed image and data analysis, MP co-conceptualized the project and supervised iMG differentation, AO co-designed image and data analysis, PS co-conceptualized and supervised the project, OB co-conceptualized and supervised the project, JHW conceptualized and supervised the project, designed experiments, co-designed image and data analysis, performed data analysis, and wrote the manuscript with input from all authors.

